# The widely used cymoxanil fungicide impairs respiration in *Saccharomyces cerevisiae* via cytochrome *c* oxidase inhibition

**DOI:** 10.1101/2024.02.29.582674

**Authors:** Filipa Mendes, Cátia Santos-Pereira, Tatiana F. Vieira, Mélanie Martins Pinto, Bruno B. Castro, Sérgio F. Sousa, Maria João Sousa, Anne Devin, Susana R. Chaves

**Affiliations:** CBMA – Centre of Molecular and Environmental Biology / ARNET – Aquatic Research Network, Department of Biology, School of Sciences, University of Minho, 4710-057 Braga, Portugal; CEB – Centre of Biological Engineering, University of Minho, Campus de Gualtar, 4710-057 Braga, Portugal; LABBELS – Associate Laboratory, Guimarães, Braga, Portugal; UCIBIO/REQUIMTE, BioSIM - Departamento de Medicina, Faculdade de Medicina da Universidade Do Porto, Alameda Prof. Hernâni Monteiro, Porto, Portugal; Associate Laboratory i4HB - Institute for Health and Bioeconomy, Faculdade de Medicina, Universidade do Porto, Porto, Portugal; CNRS, UMR 5095, Institut de Biochimie et Génétique Cellulaires, 33077 Bordeaux Cedex, France; IBS - Institute of Science and Innovation for Bio-Sustainability, School of Sciences, University of Minho, 4710-057 Braga, Portugal

**Keywords:** Cymoxanil, Mode of action, Respiration, Cytochrome *c*, Cytochrome *c* Oxidase

## Abstract

Cymoxanil (CYM) is a synthetic acetamide fungicide that has been widely used to combat downy mildew diseases in grapevine cultures and late blight diseases in tomato and potato caused by the oomycetes *Plasmopara viticola* and *Phytophthora infestans*, respectively. Despite its extensive application, the biochemical mode of action of CYM remains elusive. Previous reports indicate that CYM affects growth, DNA and RNA synthesis in *Phytophthora* and inhibits cell growth, biomass production and respiration rate in the well-characterized fungal model *Saccharomyces cerevisiae*. We therefore used this model to further characterize the effect of CYM on mitochondria. We found that CYM inhibits oxygen consumption in whole cells after 3 h of exposure, which persists over time. Using isolated mitochondria, we demonstrated that CYM specifically inhibits cytochrome *c* oxidase (C*c*O) activity during oxidative phosphorylation. Based on molecular docking algorithms, we propose that CYM acts by blocking the interaction of cytochrome *c* (cyt *c*) with C*c*O, hampering electron transfer and inhibiting C*c*O catalytic activity. Although other targets cannot be excluded, our data offer valuable insights into the mode of action of CYM that can be instrumental to drive informed management of the use of this fungicide.

## 1. Introduction

Cymoxanil [(1E)-2-(ethylcarbamoylamino)-N-methoxy-2-oxoethanimidoyl cyanide] (CYM) is a synthetic acetamide compound used as a curative and preventive fungicide. It is normally applied as a foliar spray, and is thus designated as a foliar fungicide [1]. It was first introduced in 1977, and has since been considered an important tool to protect grape and vegetable crops from oomycete pathogens of the Peronosporales order [2]. In practical applications, CYM effectively combats downy mildew diseases caused by *Plasmopara viticola* in grapevine cultures and late blight diseases triggered by *Phytophthora infestans* in tomato and potato crops [3,4]. Due to its weak persistence (half-life of a few days, depending on the conditions) and mobile nature (adsorption coefficient, K_OC_ 39–238), CYM displays a relatively short period of activity when used alone [5]. As a result, it is often combined with other fungicides, such as mancozeb and copper, which are known for their multisite mode of action and longer protective effects, to enhance overall efficacy [6].

Despite a long and extensive history of application, the precise mode of action of CYM remains unknown. According to some authors, CYM acts as a pro-fungicide and undergoes biotransformation by fungi into one (or several) fungitoxic metabolite(s). Studies in *Botrytis cinerea* demonstrated that CYM is rapidly metabolized (half-life ≤ 2 h) by a highly sensitive strain, resulting in the formation of three main metabolites, N-acetylcyanoglycine, ethylparabanic acid and demethoxylated cymoxanil. Among these, N-acetylcyanoglycine was fungitoxic against this sensitive strain, suggesting it plays an important role in the toxicity of CYM [7–9]. Mechanistically, in the case of *P. infestans*, CYM has been reported to inhibit mycelial growth and is highly effective against germ tube formation by sporangia. CYM was also shown to inhibit DNA and RNA synthesis, assessed by measuring [methyl-^3^H] thymidine and [^3^H] uridine incorporation, respectively, with DNA synthesis more sensitive to CYM than RNA synthesis. Nonetheless, no effect on RNA polymerase activity was observed, indicating that these alterations are secondary effects. Furthermore, CYM had no effect on mycelial respiration even at concentrations up to 100 mg mL^-1^ [10]. Contrasting reports can be found regarding the effect of CYM on respiration. One study reported that addition of CYM to the culture medium inhibited cell growth and biomass production and decreased respiratory rates of an *S. cerevisiae* strain [11]. Additionally, CYM inhibited ATP synthesis in a commercial *S. cerevisiae* strain [2], whereas a slight effect of CYM on the respiration of *Botrytis cinerea* was reported [12]. Beyond its primary targets (oomycetes and fungi), CYM can also affect the growth of non-target organisms, such as the bacterium *Rhodopirellula rubra* and the microalga *Raphidocelis subcapitata* [1]. These findings highlight the need for a comprehensive understanding of CYM effects on various organisms and underscore the importance of careful consideration when using this fungicide. Despite the interest from several laboratories in studying CYM cytological effects and biochemical mode of action, available data so far has not yielded conclusive results, leaving its mechanism of action still unclarified. Nonetheless, this fungicide continues to be widely employed.

In this study, we sought to investigate the role of CYM on respiration using *S. cerevisiae* cells, which have been extensively used as a eukaryotic model for molecular and cellular biology studies. Our goal was to determine whether CYM can directly affect respiration and the mitochondrial respiratory complexes, and to unveil the underlying mechanisms. After observing reduced oxygen consumption rates in *S. cerevisiae*, we isolated yeast mitochondria to identify putative mitochondrial targets of CYM. The combined analysis of these results with molecular docking data led us to propose a mechanism of respiratory impairment via cytochrome *c* oxidase (C*c*O) inhibition that has not been previously described for CYM.

## 2. Materials and Methods

### 2.1. Yeast Strain, Culture Medium, Growth Conditions and Treatment

The yeast *Saccharomyces cerevisiae* BY4742 (*Mat a; his3Δ1; leu2Δ0; lys2Δ0; ura3Δ0*) was used throughout this study. Cells were grown aerobically at 28 °C in the following medium: 0.175% yeast nitrogen base without amino acids and without ammonium sulphate (Difco), 0.2% casein hydrolysate (Merck), 0.5% (NH_4_)_2_SO4, 0.1% KH_2_PO_4_, 2% lactate (w/v) (Prolabo), pH 5.5, 20 mg L^-1^ L-tryptophan (Sigma), 40 mg L^-1^ adenine hydrochloride (Sigma), and 20 mg L^-1^ uracil (Sigma). When applicable, cells were exposed to 50 μg mL^-1^ CYM in the culture medium. Cell growth was measured indirectly as optical density at 600 nm in a Safas spectrophotometer (Monaco).

### 2.2. Mitochondria Preparation

For mitochondria preparation, yeast cells grew in YPL medium (1% yeast extract, 0.1% potassium phosphate, 0.12% ammonium sulphate, supplemented with 2% lactate as carbon source, pH 5.5) without any treatment and were harvested in stationary phase for mitochondria preparation. Yeast mitochondria were isolated from spheroplasts as described elsewhere [13] and suspended in buffer A (0.6 M mannitol, 2 mM EGTA, and 10 mM Tris-Maleate, pH 6.8). Protein Concentration was performed using the biuret method [14] using bovine serum albumin (BSA) to build the standard curve.

### 2.3. Oxygen Consumption Assays

Oxygen consumption was measured in a 1 mL thermostatically controlled chamber, at 28 °C, equipped with a Clark electrode connected to a recorder. Respiratory rates were determined from the slope of a plot of O_2_ concentration *versus* time. Respiration assays in whole cells were performed in the growth medium and, when indicated, CYM, triethyltin bromide (TET), a lipophilic inhibitor of mitochondrial ATP synthase that mimics ADP depletion [15,16], or carbonyl cyanide *m*-chlorophenylhydrazone (CCCP), a well-known uncoupling agent that increases oxygen consumption rates by dissipating the electrochemical proton gradient across the inner mitochondrial membrane, allowing the electron transport chain to function at maximum speed [17,18], were added to the chamber.

For assays with purified mitochondria, 0.2 mg of protein/ml were suspended in buffer A and ethanol (100 mM) was used as respiratory substrate. For assays performed under phosphorylating conditions (state 3, or ADP-stimulated respiration), when protons drive into the mitochondrial matrix from outside the inner membrane and ADP is consumed by phosphorylating systems, thus with oxygen consumption associated with ATP synthesis, ADP was added at the concentration indicated in the figure legends. For assays performed under non-phosphorylating conditions (state 4), when no ATP is produced and oxygen consumption is maintained at low rates [19], ADP was not added. When used, the uncoupler CCCP was used at the concentration indicated in the figure legends. For C*c*O-mediated respiration, mitochondria (0.2 mg of protein/ml) were suspended in buffer A and incubated in the presence of antimycin A (2.5 μg/mg of protein), 2.5 mM ascorbate, 100 μM *N,N,N’,N’-*tetramethyl-*p*-phenylenediamine (TMPD), and 100 mM ethanol as the respiratory substrate, as described elsewhere [20]. Schematic representation of the electron transport chain and strategy used to determine C*c*O activity is highlighted in Figure 1. All the experiments using isolated mitochondria were performed after 6 min of incubation with CYM for accurate inhibition.

**Figure 1.**
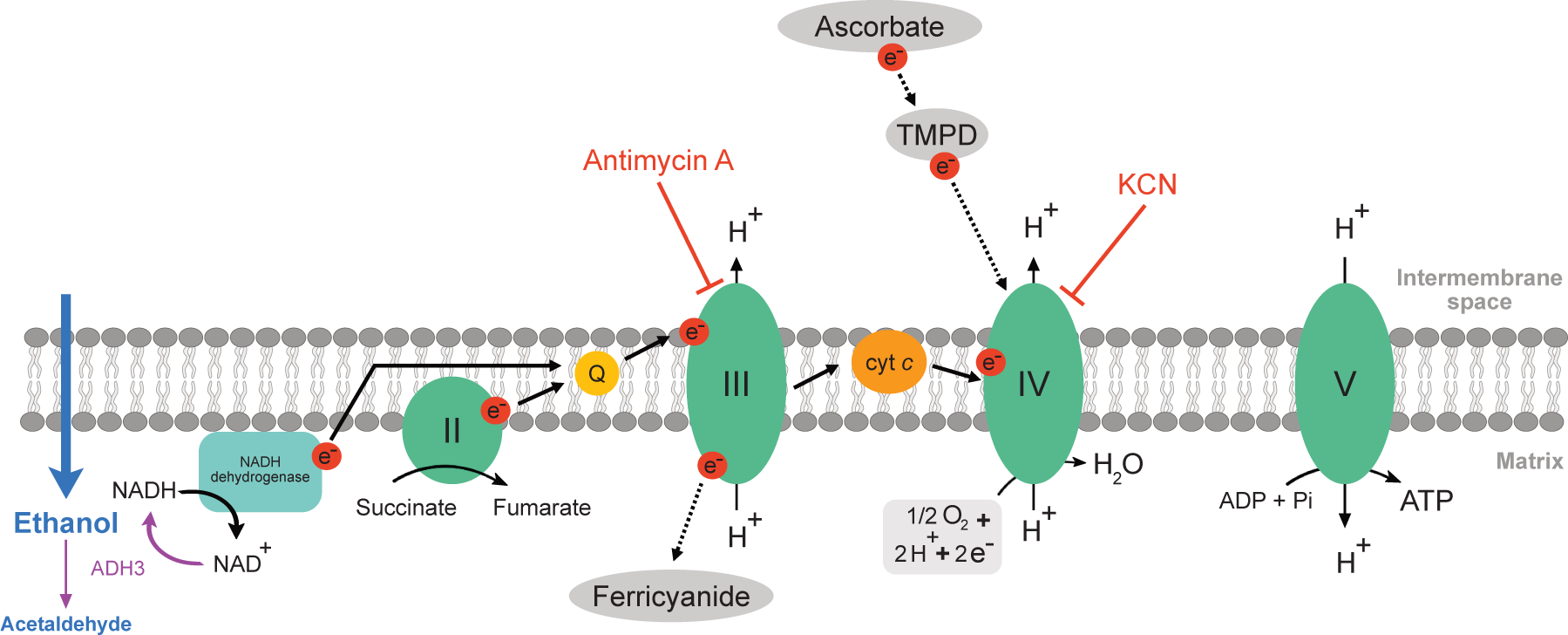
Schematic representation of the electron transport chain in *S. cerevisiae* isolated mitochondria and strategies used to determine the effect of CYM on mitochondrial complexes III and IV. All experiments were performed in the presence or absence of CYM using ethanol as substrate, thanks to the presence of mitochondrial alcohol dehydrogenase (ADH3) that catalyzes the conversion of ethanol to acetaldehyde and NADH inside the matrix [23]. The activity of complex III was based on the reduction rate of ferricyanide, an electron acceptor [21,22]. For that purpose, isolated mitochondria were incubated in respiration buffer A containing ethanol, ferricyanide and KCN (inhibitor of complex IV [24]). After monitorization of the reduction of ferricyanide, antimycin A (inhibitor of complex III [25]) was added to the reaction to ensure that the activity herein measured was indeed mediated by complex III. The activity of complex IV was based on oxygen consumption. For that purpose, isolated mitochondria were incubated in respiration buffer A with ethanol, antimycin A and ascorbate, which is the electron donor of TMPD, which in turn donates electrons to C*c*O [26,27]. After monitorization of the oxygen consumption, KCN was added to the reaction to ensure that the activity measured in this experiment was indeed mediated by complex IV.

### 2.4. Determination of Mitochondrial Complex III (Ubiquinol:Cytochrome *c* Oxidoreductase) Activity

Mitochondrial complex III activity was determined as described previously [20]. Briefly, mitochondria (0.2 mg of protein/ml) were incubated in buffer A in the presence of 1mM KCN, 1 mM ferricyanide, and 100 mM ethanol as the respiratory substrate. Ferricyanide has been used as an artificial electron acceptor, thus, under these circumstances, the ferricyanide reduction rate serves as an indicator of complex III activity [21,22]. Absorbance changes were followed at 436 nm in a Safas spectrophotometer (Monaco). The rate of ferricyanide reduction was calculated from the slope of absorbance change as a function of time. A molar extinction coefficient (ε) of 0.21 mM^-1^ cm^-1^ was used. The strategy used to determine complex III activity is systematized in Figure 1.

### 2.5. Docking Studies

To predict the binding affinity and possible modes of action of four compounds (CYM, ethylparabanic acid, N-acetylcyanoglycine and demethoxylated cymoxanil) towards C*c*O, a detailed protein-ligand docking study was performed using the GOLD software (with its four scoring functions: PLP, ChemScore, ASP and GoldScore). All the docking runs were performed in triplicate for increased sampling. The structures of the four compounds were prepared using Datawarrior [28] and Open babel [29].

Based on a deep literature search, two different hypotheses were tested to explore how CYM could potentially inhibit C*c*O: (i) docking between the heme a_3_ and the copper ion Cu_B_ at the known binding position of KCN, blocking the electron transfer mechanism. We used a cyanide ion as reference for docking coordinates and applied a 6 Å docking radius to thoroughly explore this region; (ii) docking at the interfacial region that C*c*O forms with cytochrome *c* (cyt *c)*, potentially interfering with their interaction. For this approach, the residues of chain B of C*c*O (subunit II) described by Shimada and coworkers [30] as important for protein-protein interaction with cyt *c* were used for the definition of the binding site (Ser117, Asp130, Tyr121, Asp119, Tyr105). The docking radius was 15 Å and the number of runs was set to 500, ensuring in this way that all possible configurations were considered. An additional compound, ADDA-5, (1-[2-(1-adamantyl)-ethoxy]-3-(3,4-dihydro-2(1H)-isoquinolinyl)-2-propanol hydrochloride), which was described as a C*c*O inhibitor with an IC_50_ of 21.42 µM [31], was used as a reference for the second hypothesis.

The X-ray structures selected for the preparation of the docking models were: structure 3X2Q, which contains a cyanide ion between with group heme a_3_ and Cu_B_ [24] (hypothesis 1); and structure 5IY5, which represents the X-ray structure of the C*c*O-Cyt *c* complex at 2.0-Å resolution [30] (hypothesis 2). The two structures were aligned to ensure consistency in the docking coordinates. The chains primarily evaluated in this work are highlighted in Figure 2.

**Figure 2.**
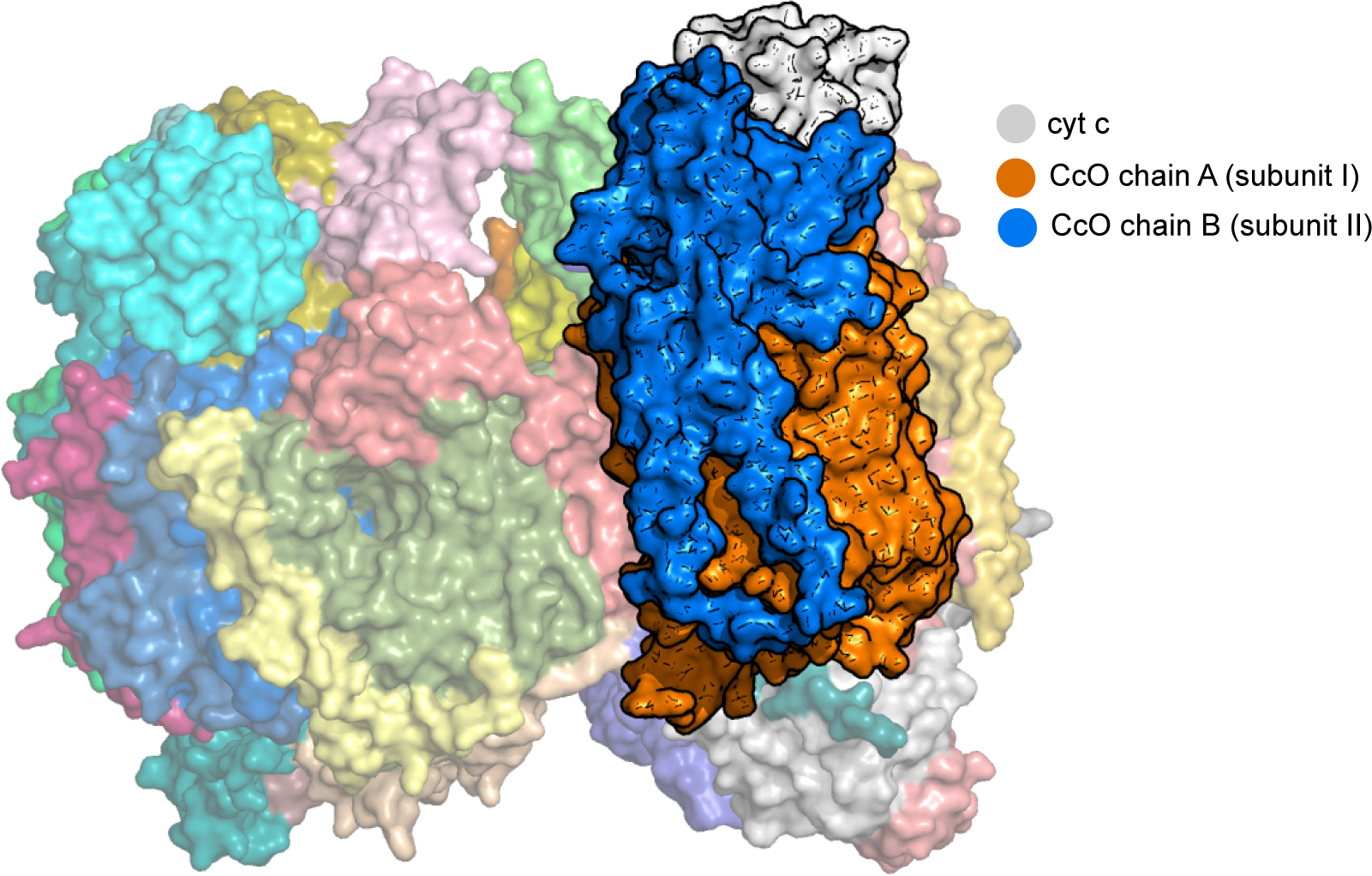
Structure of the C*c*O - cyt *c* complex used in this work. The structure (PDB: 5IY5) is shown in surface representation with all chains in a different color. The two chains evaluated for docking (in orange and blue), as well as cyt *c* (in gray) are highlighted.

### 2.6. Statistical analysis

The statistical analysis of the results was performed using GraphPad Prism software. In all assays, data are expressed as the mean and standard deviation (S.D.) of at least three independent experiments. A one-way analysis of variance (ANOVA) was used to test the effect of CYM on respiration and a Dunnett test was used to assess differences relatively to BY, state 4, state 3, state 4 + 2 μM CCCP, complex III, or complex IV. A significance level of 0.05 was employed in all analyses.

## 3. Results

### 3.1. Cymoxanil impairs respiration of whole cells

To investigate the role of CYM on respiration in whole cells, the *S. cerevisiae* BY4742 strain was first cultured in a respiratory substrate (lactic acid) with or without the addition of 50 μg mL^-1^ CYM. We found that CYM significantly inhibited the respiratory rate, reducing it by approximately 50 % after 3-4 h of treatment in comparison with non-treated cells (BY) (Figure 3). Thereafter, the percentage of inhibition remained similar until 8 h of exposure to the fungicide. Adding CYM to untreated cells directly in the electrode chamber led to a less pronounced respiration inhibition, regardless of the growth phase, although this inhibition was not statistically significant (Figure 3).

**Figure 3.**
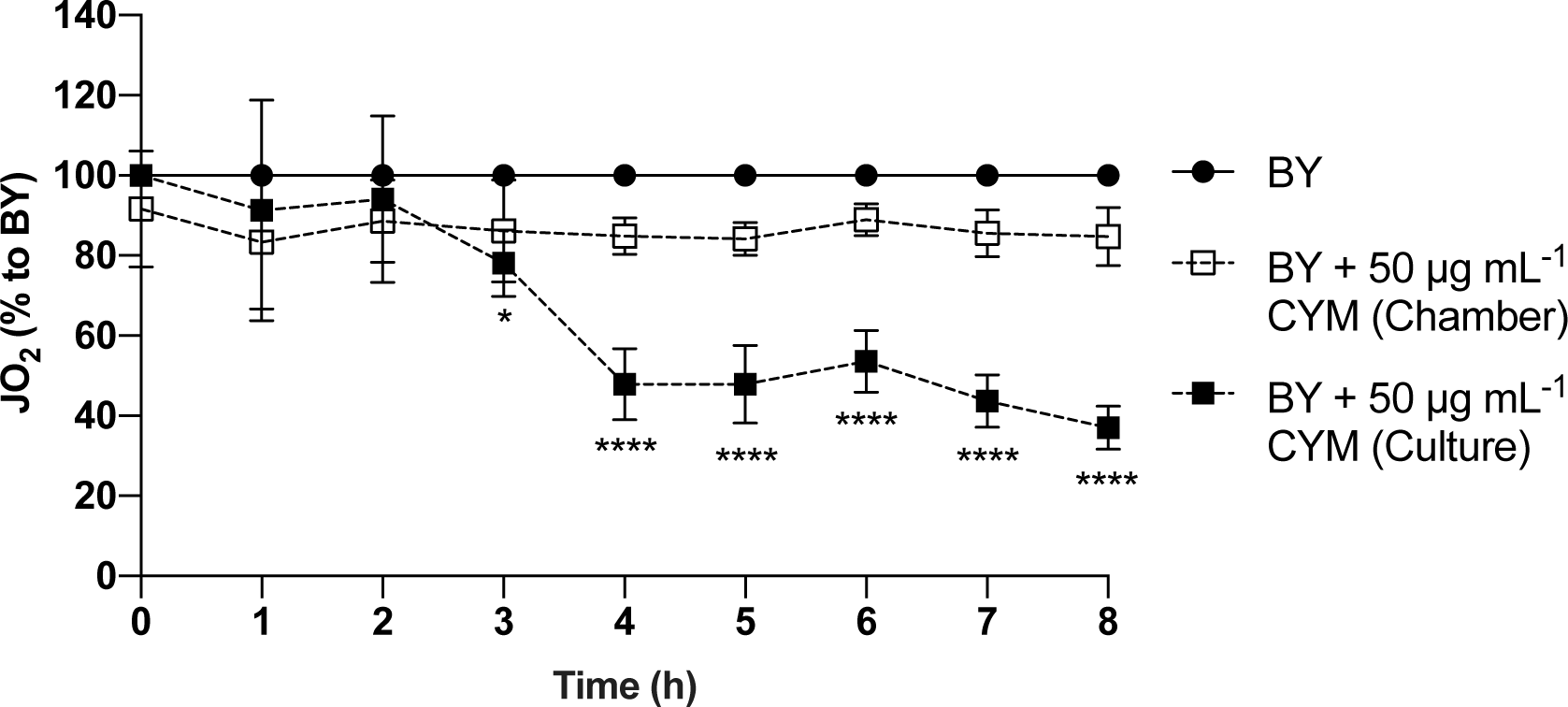
Effect of CYM on the respiratory flux (JO_2_) of yeast cells, along time. The rate of oxygen consumption of BY4742 cells was monitored as described in “Materials and Methods”. At each time point, oxygen consumption was evaluated in non-treated cells (●), cells with 50 μg mL^-1^ of CYM added in the chamber (□), and cells with 50 μg mL^-1^ of CYM added to the culture at time 0 (▪). The data are displayed as the percentage of inhibition in relation to BY (100 %) and represent mean values ± S.D. of at least three independent experiments. Asterisks (*, P ≤ 0.05; ****, P ≤ 0.0001) depict significant differences relative to BY.

We further observed that the mitochondrial ATP synthase inhibitor TET decreased respiratory rates by approximately 50 % in untreated BY4742 cells. TET also similarly decreased respiratory rates of cells grown in the presence of CYM for up to 4h, but respiratory rates decreased further in cells exposed to the fungicide for longer times (Figure 4A). As such, we observed that, after CYM inhibition of respiratory rates was apparent (from 4 h on), it could still be inhibited further by inhibition of ATP synthase.

**Figure 4.**
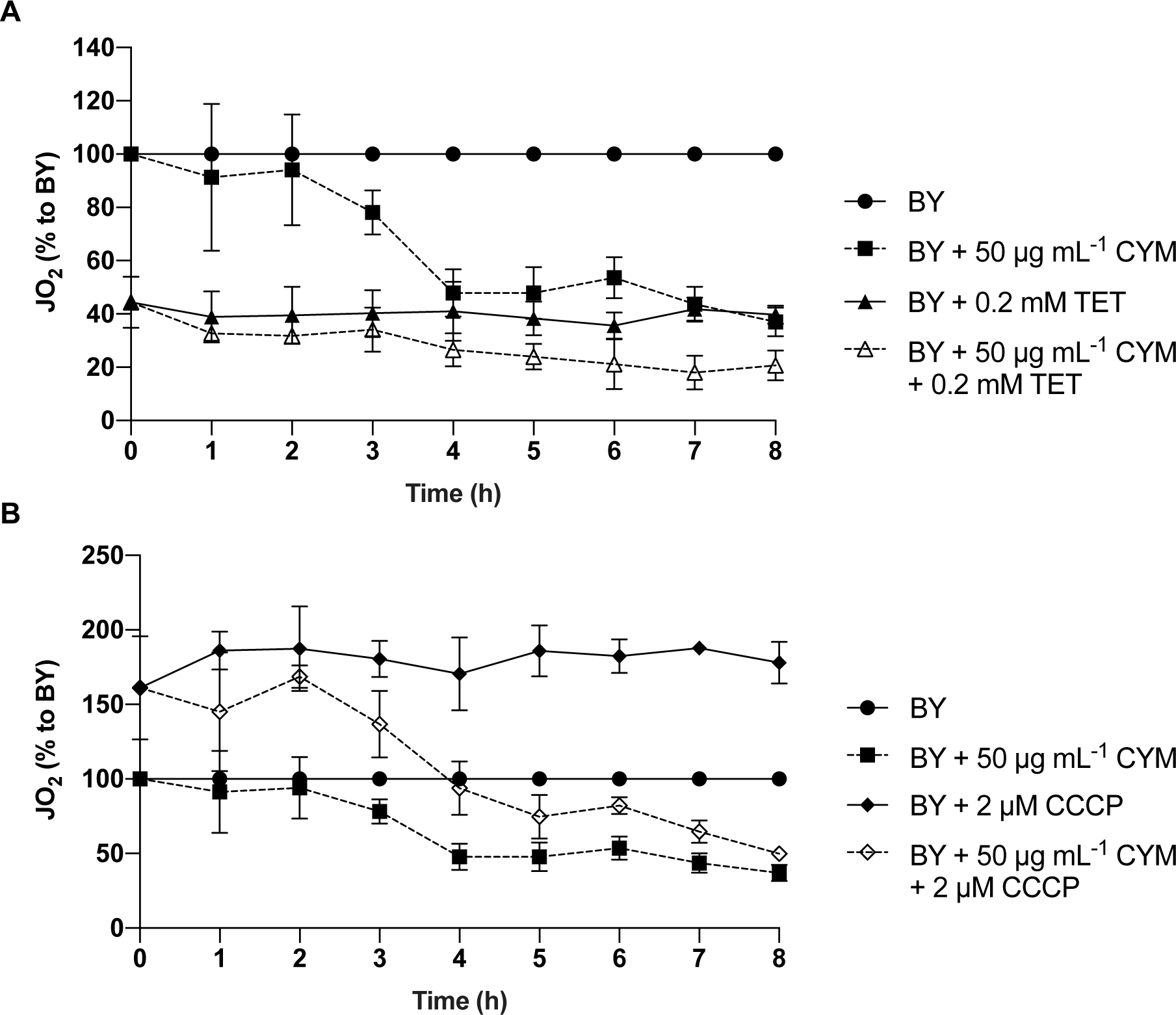
Effect of CYM on the JO_2_ of yeast cells in the presence of an inhibitor or an uncoupler. The rate of oxygen consumption was assessed as described in “Materials and Methods”, with the addition of CYM to the culture, with 0.2 mM TET (A) or 2 μM CCCP at (B) in the chamber. The data are displayed as the percentage of inhibition in relation to BY (100 %) and represent mean values ± S.D. of at least three independent experiments.

As expected, addition of the uncoupler CCCP to non-treated cells increased the respiratory flux by almost 50 % (Figure 4B). However, addition of CCCP to CYM-treated cells resulted in two different phenotypes (Figure 4B). In the first 3 h of exposure to CYM, CCCP increased the respiration rate to the maximal capacity of the mitochondrial electron transport system. However, after that, CCCP progressively lost the ability to increase respiration (Figure 4B), indicating that incubation with CYM for at least 3 h decreases the maximum respiration capacity of cells.

### 3.2. Cymoxanil compromises the mitochondrial electron transport chain

Unlike the delayed inhibition observed in whole cells, direct addition of CYM to isolated mitochondria immediately inhibited respiration (Figure 5). This inhibition was dependent on the concentration and independent of the respiratory state, as similar results were obtained under non-phosphorylating conditions (state 4, Figure 5A) and phosphorylating conditions (state 3, Figure 5B). Moreover, similarly to whole cells, we observed a concentration-dependent inhibition of state 4 respiration by CYM in the presence of CCCP (Figure 5C).

**Figure 5.**
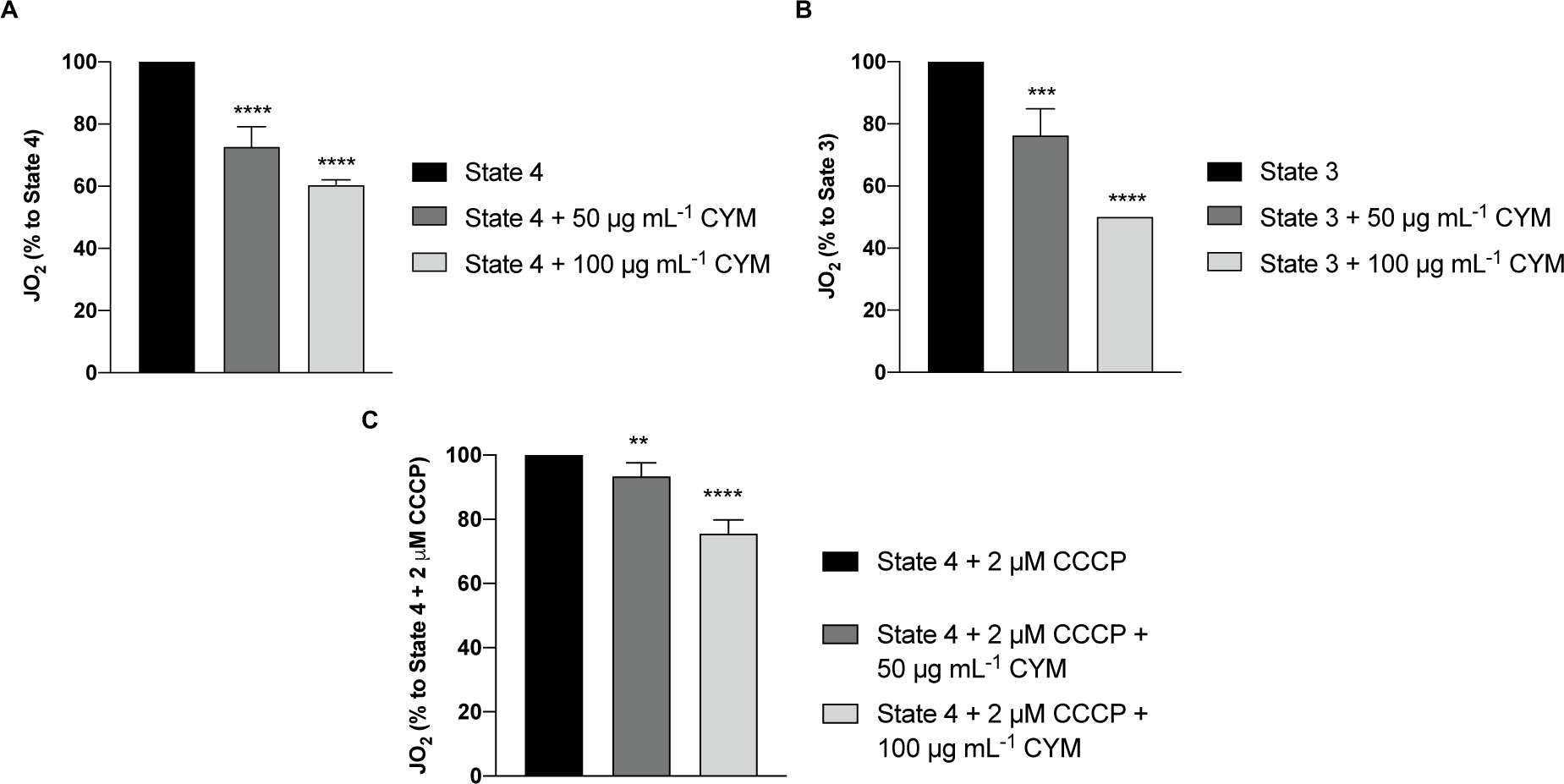
Effect of CYM on the oxygen consumption rate of isolated yeast mitochondria. Mitochondria (0.2 mg of protein/ml) were suspended in buffer A (see “Materials and Methods”) and oxygen consumption was monitored under non-phosphorylating conditions (state 4) (**A**) and phosphorylating conditions (state 3), upon addition of 1 mM ADP (**B**). The rate of oxygen consumption was also measured when state 4 was induced and 2 μM CCCP was added (**C**). All assays were performed using 100 mM ethanol as the respiratory substrate and conducted in the absence or presence of 50 and 100 μg mL^-1^ CYM. Results are expressed as a percentage of inhibition in relation to state 4, state 3 or state 4 + 2 μM CCCP (100 %) and represent mean values ± S.D. of at least three independent experiments. Asterisks (*, P ≤ 0.05; ***, P ≤ 0.001; ****, P ≤ 0.0001) depict significant differences relative to state 4, state 3 or state 4 + 2 μM CCCP.

Since we found that CYM inhibits the electron transport chain, likely independently of ATP synthase, we further assessed the effect of CYM on the activity of individual complexes. We found that complex III activity was not altered significantly, even at the higher concentration tested (Figure 6A). On the other hand, activity of complex IV (C*c*O) was strongly inhibited by both concentrations tested (Figure 6B). Moreover, the CYM effects shown in Figure 4B were sensitive to KCN (data not shown), as evidence that the oxygen consumption assessed in this assay was indeed mediated by C*c*O.

**Figure 6.**
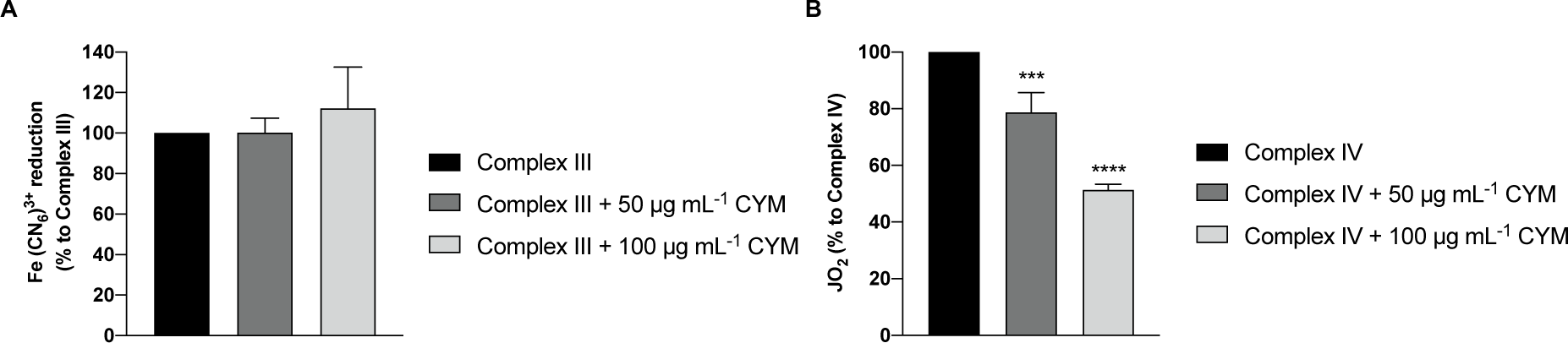
Effect of CYM on the activity of mitochondrial complex III and complex IV. **(A)** For determination of complex III activity, mitochondria (0.2 mg of protein/ml) were incubated in respiration buffer A containing 1 mM KCN, 2 mM ferricyanide and 100 mM ethanol. The reduction of ferricyanide was monitored spectrophotometrically at 436 nm, in the absence or in the presence of 50 or 100 μg mL^-1^ CYM. **(B)** For determination of complex IV (C*c*O) activity, mitochondria (0.2 mg of protein/ml) were incubated in respiration buffer A with 100 mM ethanol, 2.5 μg of antimycin A/mg of protein, 2.5 mM ascorbate, and 100 μM TMPD. Oxygen consumption was monitored in the absence or presence of 50 or 100 μg mL^-1^ CYM. Results are expressed as a percentage of inhibition in relation to complex III or complex IV (100 %) and represent mean values ± S.D. of at least three independent experiments. Asterisks (***, P ≤ 0.001; ****, P ≤ 0.0001) depict significant differences relative to complex III or complex IV.

### 3.3. Model for cymoxanil inhibition of cytochrome *c* oxidase activity

Since we found that CYM directly inhibits mitochondrial C*c*O, we sought to predict a possible mechanism by which it inhibits this complex by using computational tools. Since the respiratory rate in whole cells was significantly impaired by CYM only after 4 h incubation, and when directly added to isolated mitochondria the inhibition was less pronounced, we hypothesized that the effect of CYM on respiration of whole cells might also be linked to its metabolization and the potential action of generated metabolites. We therefore analyzed possible hypotheses of C*c*O inhibition by CYM or its metabolites: (i) docking between the heme a_3_ and the copper ion Cu_B_ at the known binding position of KCN, blocking the electron transfer mechanism; (ii) docking at the interfacial region that C*c*O forms with cyt *c*, potentially interfering with their interaction, and using ADDA-5 as a reference.

When testing the first hypothesis, we observed that the selected docking site (heme a_3_-Cu_B_ center) was strongly unfavorable, with strong steric tension, due to the very limited accessible space. Possible docking poses were obtained, but they were 5-6 Å away from this center, positioned at the surface of the protein, where the molecule interacted with the propionate groups of heme a and heme a_3_. As shown in Table S1, these values were very low, suggesting the existence of extreme steric clashes, particularly for the molecules with higher molecular weights, with demethoxylated cymoxanil and ADDA-5 exhibiting negative scores. Figure S1 depicts the docked position of each tested compound, and consistent results were observed in docking runs performed with other scoring functions such as ASP, Chemscore and GoldScore (data not shown). Further analysis of the C*c*O structure and comparison with other X-ray structures of C*c*O confirmed that there is insufficient space for molecules with more than two or three atoms to bind between the copper ion and heme a3.

The second hypothesis explored the possibility of these molecules inhibiting C*c*O activity by interfering with electron transfer between cyt *c* and C*c*O. During testing, we observed interesting and highly consistent results across different evaluation methods. Indeed, the tendency in terms of docking scores remained the same across all scoring functions tested for this hypothesis. The variation in scores is attributed to the distinct aspects of protein-ligand interactions that each scoring functions considers (Table 1).

**Table 1.** Docking results between C*c*O and the tested compound using the GOLD software. Results of the different GOLD scoring functions (PLP, ChemScore, ASP and GoldScore) for the docking between C*c*O and cymoxanil (CYM), or its potential metabolites ethylparabanic acid, N-acetylcyanoglycine and demethoxylated CYM and ADDA-5. The latter was used as a reference compound.

| COMPOUND | SCORING FUNCTIONS |  |  |  |
| --- | --- | --- | --- | --- |
|  | PLP | ChemScore | ASP | GoldScore |
| **Ethylparabanic acid** | 19.15 ± 0.01 | 14.33 ± 0.02 | 13.48 ± 0.09 | 19.69 ± 0.02 |
| **N-acetylcyanoglycine** | 24.70 ± 0.01 | 12.43 ± 0.02 | 11.78 ± 0.23 | 28.77 ± 0.56 |
| **CYM** | 31.38 ± 0.25 | 14.55 ± 0.12 | 16.40 ± 0.27 | 29.25 ± 0.07 |
| **Demethoxylated cymoxanil** | 25.06 ± 0.01 | 11.83 ± 0.20 | 15.74 ± 0.20 | 24.73 ± 0.16 |
| **ADDA-5** | 50.64 ± 0.50 | 23.91 ± 0.33 | 25.05 ± 0.81 | 40.15 ± 0.79 |

Based on the analysis of the absolute GOLD/PLP scores (Figure 7), ethylparabanic acid exhibited the lowest binding affinity, while the reference compound ADDA-5 displayed the highest affinity, primarily attributed to its much larger size and molecular weight (Figure 7). Among the test compounds, CYM presented the highest score. This consistent trend was observed across all scoring functions (Table 1).

**Figure 7.**
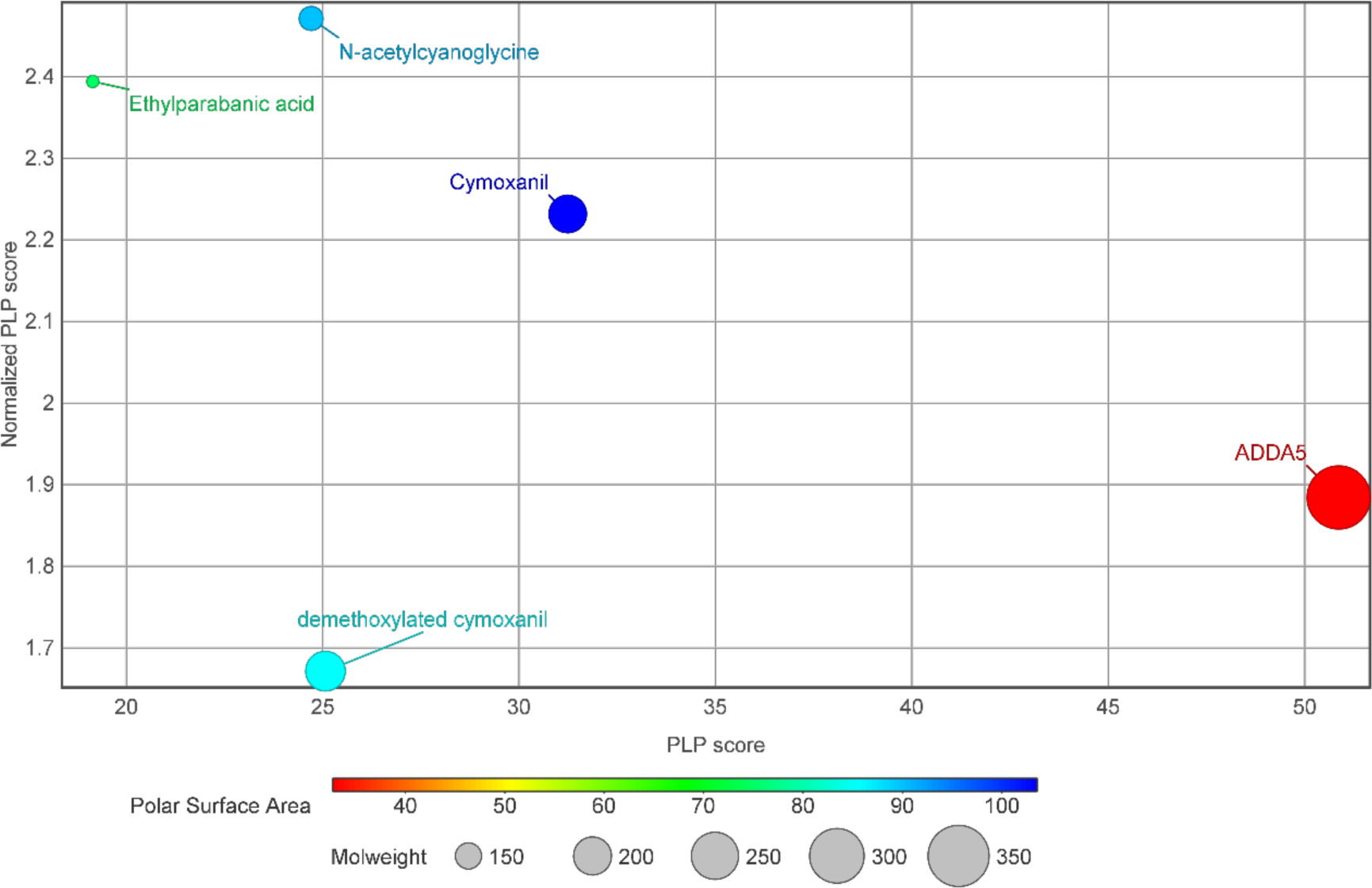
Correlation between the PLP score and the Non-H atoms of each molecule studied. The x-axis corresponds to the PLP score, while the y-axis depicts the PLP score normalized to the number of heavy atoms. The size of each marker reflects the molecular weight, and the color corresponds to the polar surface area of each compound.

To address the potential overestimation of ligands with higher molecular weight during docking, especially when comparing molecules with very different scales like the tested compounds and ADDA-5, we normalized the scoring values by the number of heavy atoms (Figure 7). N-acetylcyanoglycine and ethylparabanic acid exhibited higher normalized scores, but CYM stood out with a much higher normalized score than ADDA-5. Overall, CYM displayed both high absolute and normalized scores.

Figure 8 displays the docking poses of all the tested compounds (using the PLP scoring function) with the contacting residues (Ser117, Asp130, Tyr121, Asp119, Tyr105) highlighted in red, while also showing the typical binding region of cyt *c*. The presence of these residues suggests the possibility of an allosteric site, indicating that the tested compounds may inhibit the protein-protein interactions between cyt *c* and C*c*O, hampering electron transfer.

**Figure 8.**
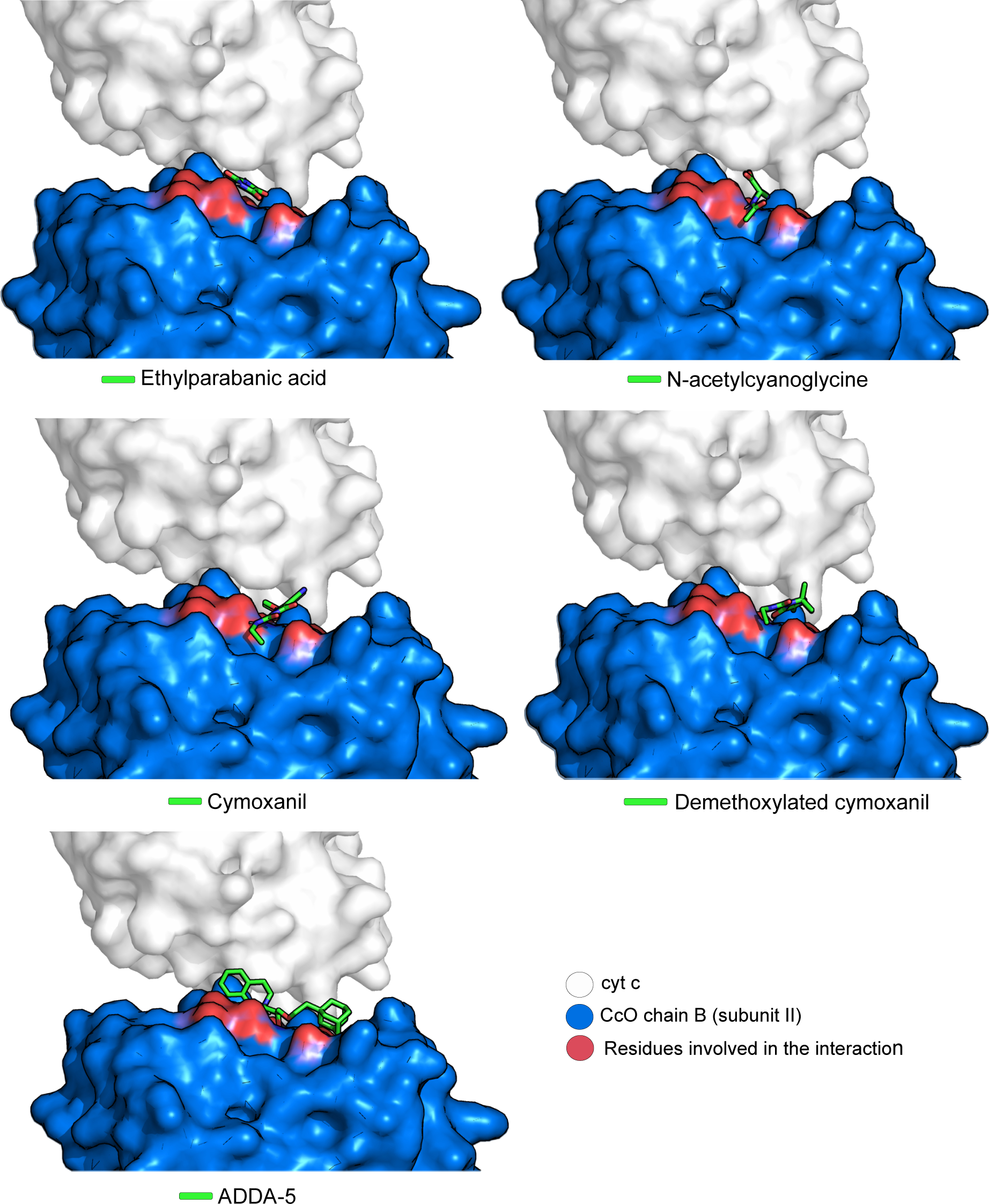
PLP docking poses for CYM, N-acetylcyanoglycine, ethylparabanic acid, demethoxylated cymoxanil and ADDA-5 on chain B (subunit II) of C*c*O. Surface representation of normal binding mode of cyt c (white) in the absence of the inhibitor molecules and of chain B of C*c*O (blue). The important residues for the interaction between cyt *c* and chain B are highlighted in red. The tested compounds are shown in sticks, with carbon atoms represented in green, oxygen atoms in red and nitrogen atoms in blue.

## 4. Discussion

In this study, we investigated the effects of CYM on oxygen consumption using a well characterized *S. cerevisiae* laboratory strain, further proposing a mechanism of action. Using whole cells, we found that inhibition of respiration by CYM was not immediate, and was mainly apparent after 4 h. Furthermore, TET decreased respiratory rates both in untreated and treated cells. However, CCCP progressively lost the ability to increase respiration in CYM-treated cells, suggesting that the proton electrochemical gradient across the membrane and the electron flow to oxygen are progressively decreased. CCCP prevents ATP synthesis by transporting protons across the mitochondrial inner membrane, interfering with the proton gradient, thus oxygen consumption of yeast cells in the presence of CCCP is independent of ATP synthase [32]. Taking into account the inability of CCCP to increase respiration of CYM-treated cells after 4 h and the fact that TET still has an effect on CYM-treated cells even after 4 h, we hypothesized that CYM acts on the mitochondrial electron transport chain. This hypothesis was supported by our findings that CYM immediately inhibited respiration of isolated mitochondria in a concentration-dependent manner under both non-phosphorylating and phosphorylating conditions, indicating that CYM inhibition is independent of the respiratory state. Altogether, these results strongly indicate that CYM directly or indirectly inhibits the electron transport chain, independently of ATP synthase.

Mitochondria of the yeast *S. cerevisiae* do not possess complex I (NADH:ubiquinone oxidoreductase) but have internal and external NADH dehydrogenases that transfer electrons to the quinone pool, thus the respiratory chain comprises three complexes: succinate dehydrogenase (complex II), complex III and C*c*O (complex IV) [33,34]. Since our results indicate that CYM does not inhibit ATP synthase, we assessed its impact on the activity of complexes III and IV. While complex III activity was not affected, CYM directly inhibited complex IV (C*c*O). As such, we sought to predict a possible mechanism using computational tools.

C*c*O is a heme/copper-containing protein with multiple subunits that plays a pivotal role in electron transfer and oxygen reduction. It receives electrons from cyt *c* and transfers them to oxygen (O_2_), reducing it to water. This electron transfer and O_2_ reduction releases free energy, which is coupled to proton pumping across the membrane [35]. While C*c*O is composed of multiple subunits, subunits I and II and their redox centers are the most important for its functioning. Subunit II contains the Cu_A_ center, which has two copper ions and acts as the initial electron acceptor from cyt *c*. It then transfers electrons to the heme a center, which in turn transmits them to the heme a_3_-Cu_B_ center, where oxygen reduction occurs. Both heme centers are located in Subunit II [24,36]. Many C*c*O inhibitors hinder its catalytic activity by targeting the heme a_3_-Cu_B_ center, including compounds like KCN [24], sodium azide [37], and nitric oxide [38]. Conversely, ADDA-5 was described as a non-competitive C*c*O inhibitor with respect to cyt *c* [31]. We therefore used computational methods to explore how CYM or three potential CYM-derived metabolites (N-acetylcyanoglycine, ethylparabanic acid and demethoxylated cymoxanil) could potentially inhibit C*c*O. Two putative mechanisms were explored: (i) docking occurs between the heme a_3_ and the copper ion Cu_B_, blocking the electron transfer mechanism (hypothesis 1); (ii) docking occurs at the interfacial region that C*c*O forms with cyt *c*, interfering with the interaction between C*c*o and cyt *c* (hypothesis 2). Values for hypothesis 1 were much lower than the typical values normally obtained in docking of reasonable inhibitors with GOLD/PLP [39]. Indeed, in the context of GOLD scoring functions, the score values are dimensionless, and a higher score is indicative of a good docking result. Furthermore, analysis of the C*c*O structure confirmed that there is insufficient space for molecules with more than two or three atoms to bind between the copper ion and heme a3, supported by the work of Yano and coworkers [15], who provided electron density maps of C*c*O bound to cyanide. Taking all these facts into account, we could confidently rule out hypothesis 1 as a plausible binding mechanism for the interaction of CYM or its metabolites with C*c*O. In contrast, highly consistent results were observed for hypothesis 2, and we therefore propose a mechanism where CYM (and its metabolites) can block the interaction of cyt *c* with C*c*O, hampering electron transfer and inhibiting C*c*O catalytic activity, by interacting with five key residues: Ser117, Asp130, Tyr121, Asp119, Tyr105. We also identified, for the first time, the potential binding site of ADDA-5, since there was previous experimental evidence that it inhibits C*c*O, but no mechanism had been proposed. Among the tested compounds, CYM exhibited a higher affinity to C*c*O than its metabolites, suggesting that CYM metabolization is not required for its inhibitory effect towards C*c*O. Our data also suggest that CYM is a promising scaffold molecule for the development of compounds designed to disrupt the C*c*O/cyt *c* interaction.

The here suggested mechanism may also be true for the target organisms of CYM commercial formulations (oomycete pathogens causers of mildews, rots, blights and other crop diseases [40]), since mitochondrial complexes and the mitochondrial respiration process are for the most part conserved [41]. Fungicides that inhibit mitochondrial respiration at complex III are highly effective against oomycetes and play an important role in downy mildew management programs [42,43]. The groups of mitochondrial complex III inhibitors that display fungicidal activity against these pathogens are quinone outside inhibitors (e.g., azoxystrobin, famoxadone and fenamidone) and quinone inside inhibitors (e.g., cyazofamid, amisulbrom). Both target the mitochondrial cytochrome *b* protein by binding at either the outer quinol-oxidizing pocket (Qo site) or the inner quinone-reducing pocket (Qi site). In both cases, the electron transfer process is blocked, leading to the inhibition of mitochondrial respiration [44–46]. Indeed, the quinone inside inhibitors used in agriculture are highly specific against oomycete pathogens [47,48]. Furthermore, it has previously been demonstrated that *P. infestans* mitochondria exhibit sensitivity to cyanide, a known inhibitor of C*c*O activity in yeast cells [49]. This indicates that are similar mechanisms between yeasts and oomycetes, suggesting that the anti-oomycete activity displayed by CYM can occur through inhibition of mitochondrial respiration by blocking the C*c*O/cyt *c* interaction, as demonstrated in this work for yeast cells. Yeast mitochondrial complexes and the mitochondrial respiration process are extremely similar to those found in higher eukaryotic organisms, including humans. Thus, this work also provides cues of the potential harmful effects of CYM in non-target organisms.

Altogether, these insights contribute to a better understanding of the mechanism of action of CYM. Further research on the specific molecular interactions between CYM and C*c*O will shed light on the basis of its inhibitory activity, aiding in the development of new antifungal agents targeting mitochondrial function.

## Abbreviations

CYM: cymoxanil
C*c*O: cytochrome *c* oxidase
cyt *c*: cytochrome *c*
CCCP: carbonyl cyanide *m*-chlorophenylhydrazone
TET: triethyltin bromide
TMPD: *N,N,N’,N’-*tetramethyl-*p*-phenylenediamine

## Acknowledgements

Investigation by the authors has been supported by national funds (Portuguese Science Foundation, FCT) via the institutional programs supporting CBMA (UIDB/04050/2020, DOI: 10.54499/UIDB/04050/2020) and ARNET (LA/P/0069/2020), and funding to Susana Chaves DOI:10.54499/DL57/2016/CP1377/CT0026. Filipa Mendes was supported by a PhD scholarship from FCT (SFRH/BD/147574/2019).

## CRediT author statement

**Filipa Mendes:** Conceptualization, Data curation, Formal analysis, Investigation, Validation, Visualization, Writing - original draft, Writing – review & editing. **Cátia Santos-Pereira** Data curation, Formal analysis, Writing - original draft, Writing – review & editing. **Tatiana F. Vieira, Mélanie Martins Pinto and Sérgio F. Sousa:** Data curation, Formal analysis. **Bruno B. Castro, Maria João Sousa, Anne Devin and Susana R. Chaves:** Conceptualization, Funding acquisition, Project administration, Resources, Supervision, Validation, Writing - original draft, Writing - review & editing. All authors read and approved the manuscript.

## Data statement

The data that support the findings of this study are available from the corresponding author, upon reasonable request.

